# CD4 T cells acquire innate capability upon classical T cell activation

**DOI:** 10.1101/2025.04.17.649269

**Authors:** Nima Yassini, Eva Goljat, Camilla Panetti, Matthias Rath, Nicole Joller

**Affiliations:** Department of Quantitative Biomedicine, University of Zurich, 8057 Zurich, Switzerland; Center for Human Immunology, University of Zurich, Zurich, Switzerland

## Abstract

Memory T cells, a sizable compartment of the mature immune system, enable enhanced responses upon re-infection with the same pathogen. We have recently shown that virus-experienced innate acting T (T_IA_) cells can modulate infectious or autoimmune diseases through TCR-independent IFN-γ production. However, how these cells arise remains unclear. Here, we show that CD4 T_IA_ cells are present in various disease settings hinting towards a disease-agnostic nature. TCR stimulation and CD28 co-stimulation are sufficient to induce naïve murine and human CD4 T cells to become capable of cytokine-mediated, TCR-independent IFN-γ responses. In true T_IA_ fashion, adoptive transfer of *in vitro*-induced T_IA_ cells in mice yielded a TCR-independent IFN-γ response during the innate phase of a *Legionella pneumophila* infection. Our data thus shows that CD4 T_IA_ cells are more ubiquitous than anticipated and could therefore be involved in more settings than expected.

## Introduction

Immunology research utilizing murine models typically uses specific-pathogen-free (SPF) mice to study various diseases where the immune system is or could be involved. This use of SPF mice has many benefits, most notably a controlled environment leading to reduced variability stemming from confounding variables and thus increased reproducibility of experimental findings. While this isolated study allows for a better mechanistic understanding of diseases, it neglects potential interactions originating from past pathogenic encounters that influence the immune composition which in turn could respond differently to subsequent heterologous challenges. This becomes especially apparent when considering the fact that in humans already at the age of 2 years, memory T cells constitute the vast majority of T cells at mucosal sites [1]. It is therefore important to also consider how past immune challenges could influence susceptibility to unrelated diseases.

We have previously shown that memory CD4 T cells generated during an acute lymphocytic choriomeningitis virus (LCMV) infection can be rapidly recruited and respond to an unrelated bacterial infection, such as *Legionella pneumophila*. Within two days, these cells produce IFN-γ through a TCR-independent mechanism, leading to a faster clearance of the pathogen [2]. This innate response could be evoked through IL-12 and IL-18 stimulation, a cytokine combination that has been shown to synergize and stimulate IFN-γ production in T cells [3–6], and to a lesser extent IL-12 and IL-33 stimulation [2]. Similar reports of human CD4 T cells have shown that they too can be activated through cytokines to produce IFN-γ [7–9].

While the murine innate acting T (T_IA_) cells were beneficial in a bacterial challenge, CD4 T_IA_ cells were detrimental to the host in an autoimmune setting as they promoted an earlier onset of experimental autoimmune encephalomyelitis (EAE) [2], an autoimmune model for multiple sclerosis. Such TCR-independent CD4 T cell-responses have also been hinted to play a role in patients suffering from autoimmunity such as rheumatoid arthritis patients [8]. These findings implicate CD4 T_IA_ cells in a range of disease contexts. However, the mechanisms underlying their generation remain unclear. In this study, we demonstrate that CD4 T_IA_ cells cells are indeed present across multiple disease settings and show that classical T cell activation alone is sufficient for both murine and human naïve CD4 T cells to gain cytokine responsiveness. We further show that, in contrast to murine CD4 T cells, human CD4 T cells are more responsive to IL-33 than IL-18 in driving cytokine-mediated, TCR-independent IFN-γ responses. Finally, *in vitro* generated CD4 T_IA_ cells were capable of responding to a *L. pneumophila* challenge in a TCR-independent manner, similar to *in vivo* generated T_IA_ cells.

## Results

### Innate acting T cells are present in diverse contexts

To gain insight into how CD4 innate acting T (T_IA_) cells are generated, we first examined in which settings CD4 T_IA_ cells can be found. To this end, mice were infected with influenza A virus or the acute WE or chronic Clone 13 (Cl13) strain of lymphocytic choriomeningitis virus (LCMV). At least 40 days after the acute infections (WE memory or Flu memory) or 25 days into the chronic infection (Cl13 chronic), we examined the CD4 T cells found in the lung of these mice. Using the CD4 T_IA_ markers IL-18R and CXCR6 [2], we could clearly observe an increased expression compared to naïve controls (Figure 1A). We then further analyzed this population using unsupervised clustering and observed a dominant cluster 2 with higher VLA-4 expression in WE memory (Figure 1B–D), as we have previously shown [2]. While Flu memory mice featured a prominent cluster 8 marked by higher CXCR3 expression, and mice chronically infected with LCMV Cl13 displayed a dominant cluster 3 with high PD-1 expression. Nevertheless, all clusters were represented in each of the infectious settings, suggesting that CD4 T_IA_ cells capable of responding to cytokine stimulation alone, could be generated in divers infectious setting, although with alterations in terms of proportion and expression of additional markers.

**Figure 1.**
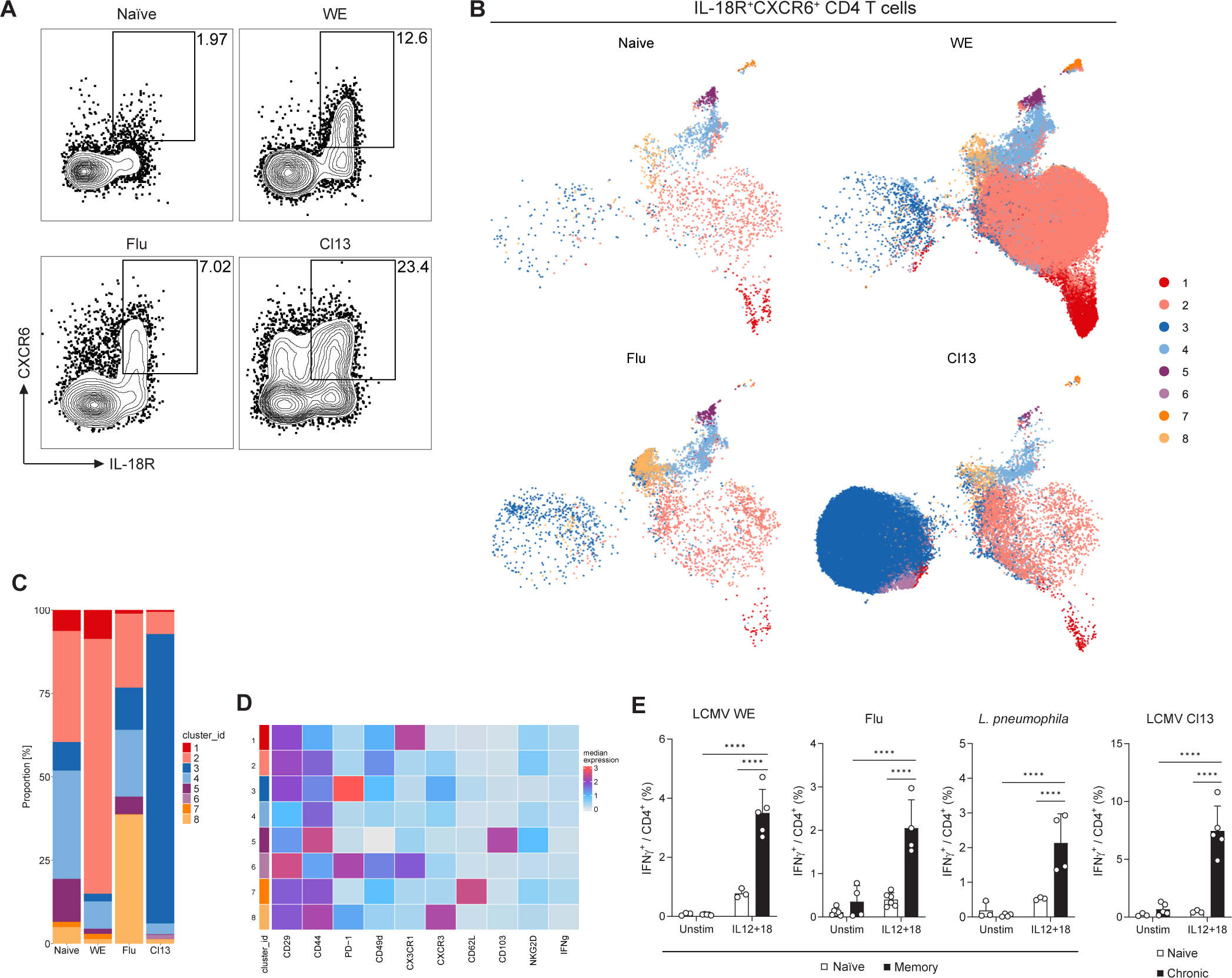
CD4 T_IA_ cells found after various infections. (A–D) Mice were infected with LCMV WE (i.v.), LCMV Cl13 (i.v.), influenza A virus (Flu, i.n.) or left naïve and immune cells of the lung were isolated from LCMV WE or Flu infected mice (day ≥ 40) or LCMV Cl13 infected mice (day ≥ 25) and stained for IFN-γ production. (A) Representative FACS plots gated on CD4 T cells. (B–D) IL-18R^+^CXCR6^+^ CD4 T cells, gated as shown in (A), were further analyzed using unsupervised clustering (n=2–5). (B) UMAP, (C) proportion of each cluster per condition and (D) heatmap showing marker expression within these clusters are depicted. (E) Splenic CD4 T cells of LCMV WE, *L. pneumophila*, and LCMV Cl13 experienced mice and lung CD4 T cells of Flu-experienced mice were isolated and stimulated overnight with cytokines or left untreated (n=3–6, two-way ANOVA with Šídák).

We therefore asked if these cells, despite their differences in marker expression (VLA-4, CXCR3, PD-1), indeed possess the functional innate capability to mount an IFN-γ response upon cytokine stimulation. To this end, we isolated CD4 T cells form mice that had undergone and cleared infection with LCMV WE, influenza A, or *L. pneumophila* or were chronically infected with LCMV Cl13. CD4 T cells were then stimulated with IL-12 and IL-18, which have been shown to be relevant for the T_IA_ response [2]. As previously described, LCMV WE memory CD4 T cells showed a significantly higher IFN-γ production compared to the naïve controls (Figure 1E). Besides this acute systemic viral infection, the acute local lung infection with influenza, the acute systemic bacterial *L. pneumophila* infection, and even the chronic systemic LCMV Cl13 infection all induced a CD4 T cell population equipped with enhanced responsiveness to cytokine stimulation leading to IFN-γ production.

Taken together, these results suggest that the innate acting capability of Th1 cells could be more ubiquitous than anticipated, as their responsiveness to cytokine stimulation alone could be detected in a broad range of infections.

### Human CD4 T_IA_ cells are more responsive to IL-33 than IL-18

Because of the diverse settings in which CD4 T_IA_ cells are present, we next examined human CD4 T cells for their TCR-independent cytokine responsiveness. Previous reports have shown that IL-12 and IL-18 in combination with some common gamma chain cytokines can elicit an IFN-γ response in human CD4 T cells [7–9]. We have previously demonstrated that IL-12 and IL-33 can also elicit an IFN-γ response in murine CD4 T cells, although to a lesser extent than IL-12 and IL-18 [2]. However, whether IL-33 can activate human CD4 T cells, has not been explored. To this end, we isolated CD4 T cells from PBMCs of healthy donors, sorted CD45RA^+^CD45RO^−^ CD4 T cells (naïve) and CD45RA^−^CD45RO^+^CXCR3^+^ CD4 T cells (memory Th1) and stimulated them with combinations of IL-12 plus IL-18 or IL-33, in the presence or absence of IL-2 and IL-15 (Figure 2A). As previously reported [8], the combination of IL-12 and IL-18 together with IL-15 or with IL-2 and IL-15 was able to evoke an IFN-γ response. To our surprise, the combination of IL-12 and IL-33 together with IL-2 and/or IL-15 lead to an even greater IFN-γ response than that induced by IL-18.

**Figure 2.**
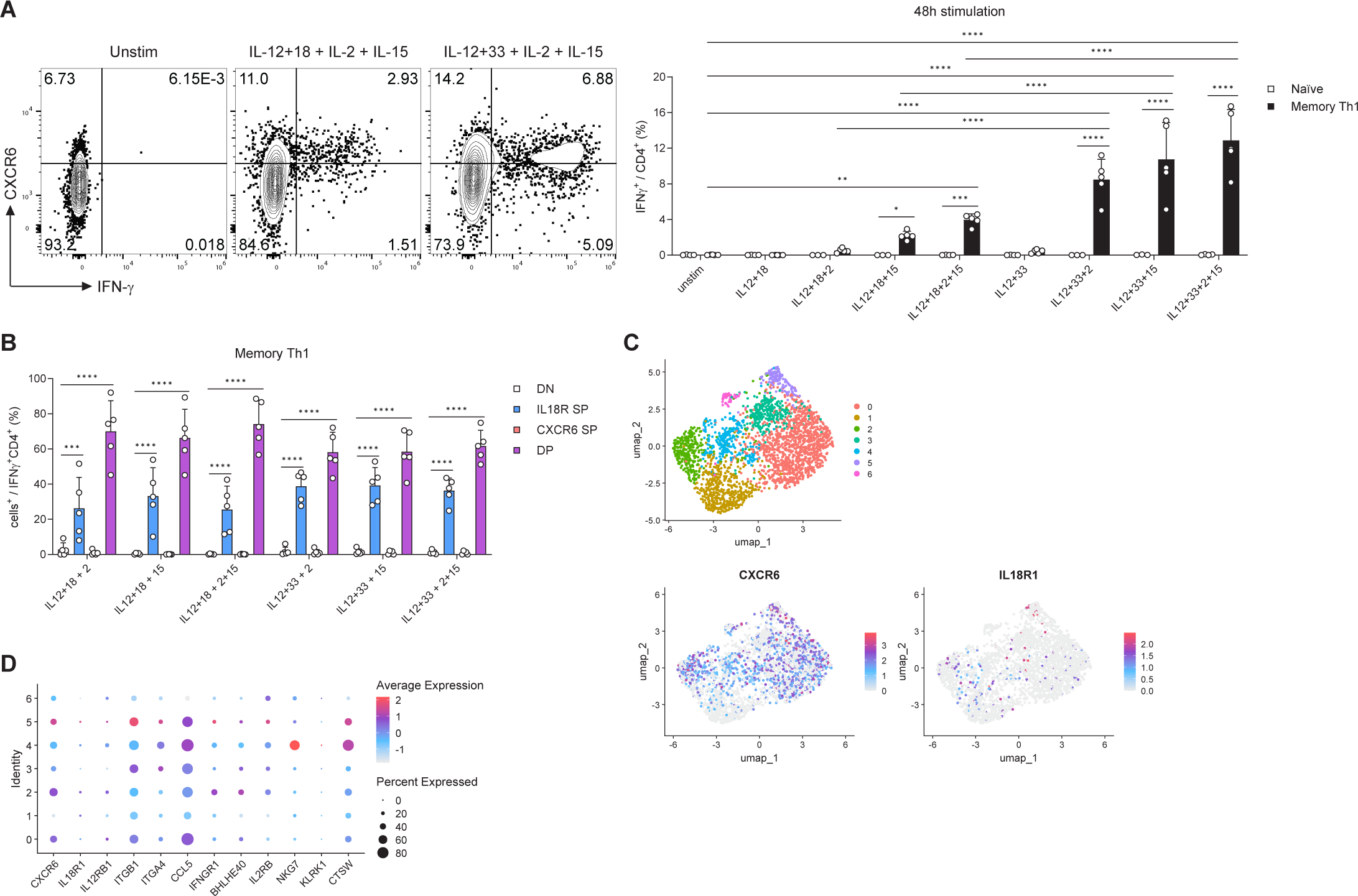
Human CD4 T cells respond strongly to IL-12, IL-33, IL-2 and/or IL-15. (A and B) Human CD4 T cells isolated from PBMCs of healthy donors were sorted for CD45RA^+^CD45RO^−^ (Naïve) or CD45RA^−^CD45RO^+^CXCR3^+^ (Memory Th1) and stimulated with the indicated cytokines (n=4–5). (A) The resulting IFN-γ response is shown in representative FACS plots (left, gated on CD4 T cells) and summary graphs (right; two-way ANOVA with Šídák). (B) Frequency of marker^+^ populations among IFN-γ^+^ CD4 T cells (two-way ANOVA with Dunnett). (C and D) Single-cell RNAseq dataset of human lung CD4 T cells [10] analyzed for T_IA_ marker expression displayed as (C) UMAP or (D) Dotplot.

Looking at the expression of IL-18R and CXCR6, a combination of receptors previously identified as an effective marker combination for murine CD4 T_IA_ cells [2], we observed that most of the IFN-γ producing cells co-expressed IL-18R and CXCR6 (Figure 2B), similar to the murine setting. However, a significant portion of the IFN-γ producing human cells was also single-positive for IL-18R, suggesting that for human CD4 T cells IL-18R expression alone serves as a better marker for potential CD4 T_IA_ cells. These differences between murine and human CD4 T_IA_ cells prompted us to analyze expression of T_IA_ signature genes (derived from their murine transcriptional profile [2]) in a dataset of human CD4 T cells from healthy lung tissues [10] (Figures 2C & D). While some CD4 T_IA_ genes were barely detectable or expressed to a lower extent in human CD4 T cells (*IL18R1, IL12RB1*, *ITGA4*, *IFNGR1*, *BHLHE40*, *IL2RB*, *KLRK1*), others (*CXCR6*, *ITGB1*, *CCL5*, *NKG7*, *CTSW*) were more highly expressed. Together, these findings show that, while functional CD4 T_IA_ cells are present in humans, these cells phenotypically differ from their murine counterpart. Most notably, human CD4 T_IA_ cells show a significantly greater responsiveness to IL-33 compared to IL-18 when combined with IL-12, IL-2 and/or IL-15.

### *In vitro*-generated CD4 T_IA_ cells are functional *in vivo*

Given that the cytokine responsiveness could be detected in both murine and human CD4 T cells and the diverse disease settings from which murine CD4 T_IA_ cells can originate, we sought to determine when during the infection they first appear. We infected mice with LCMV WE and isolated CD4 T cells at days 4, 5, 10, 15, 25, and 40 of the infection, which were then stimulated with our cytokine combination to determine their ability to produce IFN-γ in response to this TCR independent stimulation. To our surprise, the responsiveness kinetic of the CD4 T cells mirrored the classical T cell activation kinetic with a peak response at day 10, followed by a contraction and a memory phase (Figure 3A).

**Figure 3.**
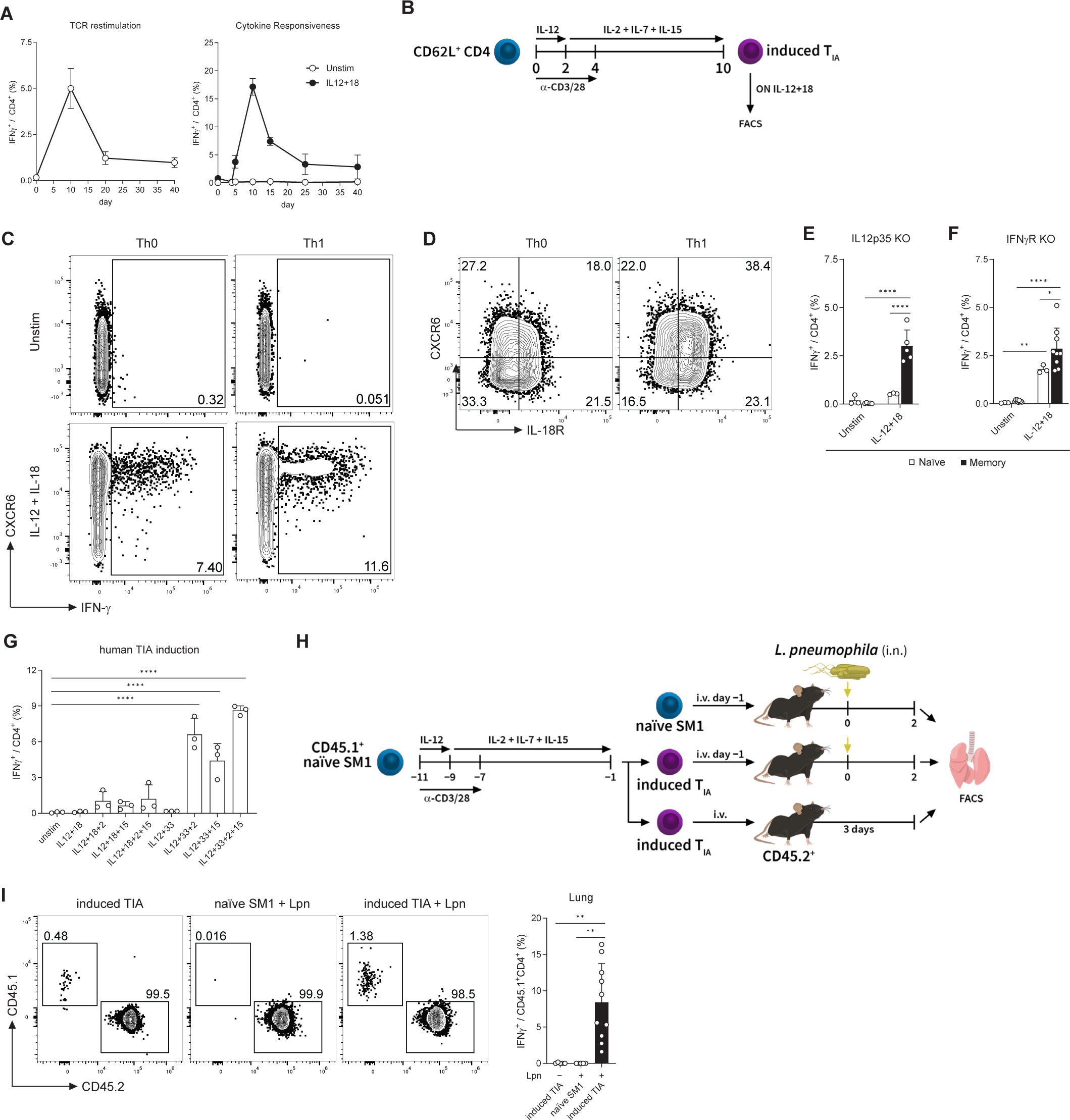
CD4 T_IA_ cells can be induced *in vitro* and are functional *in vivo.* (A) CD4 T cells of LCMV-experienced mice were isolated at indicated time points and re-stimulated with gp61 (left) or stimulated overnight with cytokines or left untreated (right) and analyzed for IFN-γ production. (B) Experimental scheme of the induction protocol used in (C) and (D). Th0 cells were not polarized by IL-12 in the first two days of the protocol. (C) IFN-γ response of CD4 T cells after completing the induction protocol. (D) IL-18R and CXCR6 expression of CD4 T cells after the 10-day protocol without additional stimulation. (E and F) Splenic CD4 T cells of naïve or LCMV WE memory mice were isolated and stimulated overnight with cytokines or left untreated. KO mice for IL-12p35 (E) and IFNγR (F) were tested (D: n=3–5; E: n=3–9; two-way ANOVA with Šídák). (G) Sorted CD45RA^+^CD45RO^−^ human CD4 T cells induced using the protocol depicted in (B) were stimulated with cytokines for 48h to determine their IFN-γ response (n=3, one-way ANOVA with Dunnett). (H) Experimental setup used in (I). (I) Representative FACS plots gated on CD4 T cells (left) and IFN-γ response of SM1 cells (right) 3 days after adoptive transfer (n=5–10; one-way ANOVA with Kruskal-Wallis; Lpn: *L. pneumophila*).

Given the similarities of the kinetics of CD4 T_IA_ responsiveness and the classical T cell response, we set out to determine whether CD4 T_IA_ cells could be induced by classical T cell differentiation of naïve CD4 T cells *in vitro*. We sorted splenic CD62L^+^ CD4 T cells from naïve mice and stimulated them for 4 days with anti-CD3 and anti-CD28 antibodies followed by a 6-day resting period (Figure 3B). IL-12 was added during the first 2 days of the protocol to polarize cells towards Th1, while cells without IL-12 (Th0) were used as controls. After this 10-day protocol, cells were stimulated with IL-12 and IL-18 overnight or left unstimulated. Surprisingly, not only *in vitro* generated Th1 cells, but also Th0 cells could respond to cytokine stimulation and produce IFN-γ (Figure 3C), revealing that classical activation through TCR together with co-stimulation is sufficient to induce CD4 T_IA_ cells. Nevertheless, the Th1 polarization through addition of IL-12 during the first 2 days of the protocol yielded a greater frequency of responding cells capable of producing IFN-γ. Additionally, the Th1 cultures contained an increased population of cells co-expressing the CD4 T_IA_ markers IL-18R and CXCR6 (Figure 3D). As IL-12 and IFN-γ both play a role in the polarization of Th1 cells, we explored their requirement for the generation of CD4 T_IA_ cells. Mice deficient in IL-12 (IL12p35 KO) or lacking the IFN-γ receptor (IFNγR KO) were infected with LCMV WE and allowed to clear the infection and develop T cell memory (at least 40 days post infection). Cytokine stimulation of CD4 T cells isolated from naïve and memory mice revealed that neither IL-12 nor IFN-γ signaling are essential for CD4 T_IA_ cell generation as CD4 T cells from both memory IL12p35 KO and memory IFNγR KO showed higher IFN-γ response upon IL-12 and IL-18 stimulation than naïve controls (Figures 3E & F). Taken together, these results indicate that Th1 polarizing cytokines are not essential for the generation of CD4 T_IA_ cells but can augment the proportion of CD4 T cells that gain T_IA_ functionality.

Next, we explored whether *in vitro* differentiation of naïve human CD4 T cells could render them responsive to cytokine stimulation. For this, sorted CD45RA^+^CD45RO^−^ CD4 T cells were differentiated using the same induction protocol as for generating murine CD4 T_IA_ cells *in vitro* (Figure 3B). Indeed, after *in vitro* differentiation of naïve CD4 T cells, cytokine stimulation with IL-2 and/or IL-15 in combination with IL-12 and IL-33, but not IL-12 and IL-18, was able to elicit an IFN-γ response (Figure 3G). Thus, similar to murine CD4 T cells, human CD4 T cells can be differentiated *in vitro* to become responsive to cytokine stimulation, and, like the memory Th1 sorted CD4 T cells from healthy donors (Figure 2A), *in vitro*-generated CD4 T_IA_ cells respond more strongly to IL-33 than IL-18 stimulation. While the *in vitro*-generated CD4 T_IA_ cells phenotypically and functionally resemble those generated *in vivo*, it is unclear whether these cells also have the same *in vivo* functionality. To test this, we induced CD4 T_IA_ cells from naïve Smarta (SM1) mice, which bear a transgenic TCR specific for the LCMV gp61 peptide but do not recognize *L. pneumophia* [2, 11]. To this end, naïve or *in vitro*-generated SM1 CD4 T_IA_ cells were transferred into congenically-marked naïve recipient mice which were then infected intranasally with *L. pneumophila* or left naïve as controls (Figure 3H). On day 2 of the infection, lungs were harvested and analyzed for the IFN-γ response. Importantly, cells were only incubated with Brefeldin A and not re-stimulated, in order to determine whether a bona fide T_IA_ response is occurring. Mice that had received T_IA_-induced SM1 cells showed a higher frequency of transferred cells compared to mice that had received naïve SM1 cells (Figure 3I). While transferred cells could be detected in the lungs of both groups that received T_IA_-induced SM1 cells, an IFN-γ response could only be observed upon *L. pneumophila* infection. This TCR-independent response thus confirms that *in vitro*-induced T_IA_ cells can indeed act in an innate manner *in vivo*.

Thus, classical TCR stimulation and CD28 co-stimulation is sufficient for the *in vitro* generation of CD4 T_IA_ cells from naïve CD4 T cells, but it can be further enhanced by IL-12-mediated polarization. Importantly, *in vitro*-induced CD4 T_IA_ cells are functional *in vivo* and respond in an innate manner upon infectious challenge.

## Discussion

Unlike controlled experimental conditions in murine models, individuals in the general population are continuously exposed to a wide range of immunological challenges. Through the generation of immunological memory, this constant environmental interface shapes the immune system and may contribute to the increased inter-individual variability observed with age [12]. We have previously demonstrated that past encounters can modulate the outcome of heterologous challenges through CD4 T_IA_ cells [2]. Here we show that Th1 cells generated in diverse settings possess the capacity to respond to cytokine stimulation in a TCR-independent manner. Upon classical T cell activation, Th1 cells gain the ability to produce IFN-γ in response to cytokine stimulation alone. This innate-like function was also obtained by *in vitro*-generated T_IA_ cells, which were able to exert TCR-independent responses *in vivo* following *L. pneumophila* challenge.

In our previous work, we identified IL-18R and CXCR6 co-expression as a prominent marker combination for CD4 T_IA_ cells [2]. Here, we show that IL-18RLCXCR6L CD4 T cells are present following acute or chronic LCMV infection, as well as after local influenza infection. CD4 T cells isolated after clearance of the acute infections or during the chronic phase of LCMV Cl13 infection were able to produce IFN-γ in response to IL-12 and IL-18, regardless of the specific pathogen encountered. This cytokine responsiveness is detectable during the acute, contraction, and memory phases of infection and can be induced *in vitro* through TCR and co-stimulatory signals alone. The consistent emergence of CD4 T_IA_ cells across multiple Th1-dominated infections suggests that this acquired cytokine responsiveness and innate capability is a common feature of CD4 T cell differentiation. Indeed, this TCR-independent function is not restricted to Th1 cells, as Th2 and Th17 cells have also been shown to respond to cytokine stimulation or heterologous challenge by producing IL-13 and IL-17A, respectively [4, 13, 14].

Similar to murine CD4 T_IA_ cells, human memory Th1 cells from healthy donors were able to produce IFN-γ in response to cytokine stimulation. While murine CD4 T cells respond more robustly to IL-12 + IL-18 than to IL-12 + IL-33 stimulation [2], human CD4 T cells responded most effectively when IL-33 was used in combination with IL-12, IL-2, and IL-15. This was particularly pronounced in CD4 T_IA_ cells differentiated *in vitro* from naïve CD4 T cells.

Importantly, *in vitro* generated CD4 T_IA_ cells are functional *in vivo*, and produce IFN-γ in an antigen-independent manner during the early, innate phase of an unrelated infection. Given the minimal requirements for naïve CD4 T cells to become receptive to cytokine stimulation, it stands to reason that CD4 T_IA_ cells are likely more ubiquitous than previously appreciated. Their capacity to modulate heterologous immune responses *in vivo* - demonstrated by their protective role during infection and their contribution to the accelerated onset of autoimmunity [2] - suggests that CD4 T_IA_ cells may play a broader role across various disease settings and thus may represent interesting targets for therapeutic intervention. For instance, a recent study found that a majority of CD4 T cells infiltrating lung and colorectal tumors are likely tumor-unspecific bystander cells [15]. Targeted cytokine delivery to the tumor microenvironment could provide a means to activate these bystander cells and induce effector responses at this site. Currently, strategies are being developed to deliver cytokines such as IL-2, IL-12 and IL-18 to the tumor microenvironment [16–18]. However, our data suggest that a more effective approach may involve the combination of IL-12 and IL-33, along with CD122-biased IL-2 [18].

In summary, our findings demonstrate that classical T cell activation via TCR and CD28 co-stimulation is sufficient for naïve CD4 T cells to acquire innate-like functionality *in vitro*. Cytokine stimulation alone is sufficient to trigger IFN-γ production in CD4 T_IA_ cells, whether generated *in vitro*, isolated from murine infection models or human PBMCs. These results reveal a broadly adopted, innate-acting capability within activated and memory CD4 T cells, with potential implications for both infection and immunotherapy.

## Materials and Methods

### Mice

C57BL/6 (B6) mice were purchased from Janvier Labs and IL12p35 KO (Jackson Laboratory #002692, [19]), IFNγR KO (Jackson Laboratory #003288, [20]), and Smarta [11] mice have been previously described. Animals were bred and housed in SPF facilities at LASC Zürich, Switzerland. All experiments were performed in accordance with institutional policies and regulations of the relevant animal welfare acts and have been reviewed and approved by the Cantonal veterinary office.

### Viral and bacterial infections and adoptive cell transfers

For the acute and chronic LCMV infections, 200 FFU LCMV WE and 2×10^6^ FFU LCMV Cl13, respectively, were injected i.v.. Influenza A virus infections were achieved with 200 PFU of influenza PR8-gp33 virus given intranasally. For systemic *L. pneumphila* infections, 3×10^6^ CFU of *L. pneumphila* JR32 FlaA^−^ [21] were injected intravenously, 37 days later a booster of 3×10^6^ CFU (i.v.) was given and after at least 40 days after the booster mice were analyzed. For local *L. pneumphila* infections, 3×10^6^ CFU of *L. pneumphila* JR32 FlaA^−^ was given intranasally.

For SM1 cell transfers, CD4 T cells from SM1 mice (CD45.1) were purified using MojoSort Mouse CD4 Nanobeads (BioLegend) and 2.5×10^5^ naïve or T_IA_-induced SM1 cells were adoptively transferred i.v. into B6 recipient mice one day prior to local *L. pneumphila* infection.

### In vitro assays

To obtain single-cell suspensions, spleens were mechanically disrupted and red blood cells were lysed with ACK buffer (155 mM NH4Cl, 10mM KHCO3, 0.1 mM Na2EDTA, pH: 7.4) for 3 min. Lungs were enzymatically digested with collagenase D (Gibco) and DNase I (VWR) for 30 min, and immune cells isolated using a 30% Percoll (GE Healthcare) gradient. Human blood was collected from healthy donors, in accordance with the approved study by the Cantonal Ethics Committee of Zurich (BASEC 2024-01233). PBMCs were isolated using a Ficoll gradient (Cytiva).

For the isolation of CD4 T cells, cells were magnetically sorted using MojoSort Mouse CD4 Nanobeads (BioLegend) or human CD4+ T Cell Isolation Kit (Miltenyi Biotec). Upon isolation, 2×10^5^ murine CD4 T cells were incubated with IL-12 and IL-18 (both at 10 ng/ml) for 12–18h overnight in U-bottom plates or re-stimulated for 4h with gp61-80 (gp61, 1μg/ml, GLNGPDIYKGVYQFKSVEFD, EMC microcolletions GmbH). Human CD4 T cells were further FACS sorted into naïve and memory Th1 cells and 2×10^5^ cells were stimulated for 48h with IL-2 (100U/ml), IL-12 (10ng/ml), IL-15 (20ng/ml), IL-18 (50ng/ml) or IL-33 (50ng/ml) in U-bottom plates.

For *in vitro* induction of T_IA_ cells, 10^5^ naïve CD4 T cells were stimulated in flat bottom plates coated with anti-CD3 (mouse: 2 μg/ml, clone 145-2C11, BioXCell; human: 3μg/ml, clone UCHT, BD Biosciences) in the presence of soluble anti-CD28 (mouse: 2 μg/ml, clone PV-1, BioXCell; human: 1μg/ml, clone 28.2, BD Biosciences) for 96h in RPMI 1640 media (Gibco) supplemented with 10% heat-inactivated FCS, sodium pyruvate (Gibco), 200μM L-Glutamine (Gibco), 10U/ml Penicillin-Streptomycin (Gibco), MEM non-essential amino acids (Gibco), MEM vitamin solution (Gibco), and 50μM β-mercaptoethanol. Where indicated, cells were additionally incubated with IL-12 (10ng/ml) during the first 48h of the stimulation. From the third day onwards, cells were incubated with survival cytokines IL-2 (mouse: 20U/ml; human: 100U/ml), IL-7 (10ng/ml), and IL-15 (mouse: 50ng/ml, human: 20ng/ml). After the 96h of stimulation, cells were switched to U-bottom plates for the remaining 6 days of rest and subsequent cytokine stimulations. Recombinant cytokines were purchased from Biolegend with the exception of human IL-2 (R&D Systems). All incubations were done at 37°C and 10% CO_2_.

### Flow cytometry

FACS stainings were performed on single-cell suspensions. When staining for intracellular cytokines, cells were incubated with Brefeldin A (BioLegend) for 4h at 37 °C in 10% CO_2_ prior to staining. Fluorescently labelled antibodies were purchased from BD Bioscience, invitrogen, and BioLegend. Zombie NIR fixable dye (Biolegend) or LIVE/DEAD Fixable Blue dye (invtirogen) was used to exclude dead cells and debris. Cells were permeabilized and fixed using the Cytofix/Cytoperm kit (BD Biosciences).

BD FACSAria III or BD FACSymphony S6 was used for cell sorting. Data was acquired on a BD LSR Fortessa, BD FACSymphony A5 analyzer (BD Biosciences), or Cytek Aurora and analysed using SpectroFlo (Cytek) and Flowjo software (TreeStar).

### Analysis of flow cytometry and single-cell RNAseq data in R

For the analysis of flow cytometry data in R, FlowJo gating was used to select the population of interest by using the packages CytoML [22], flowCore [23], and flowWorkspace [24]. Dimensionality reduction and visualization was done using the CATALYST package [25, 26].

The single-cell RNA-seq dataset from Travaglini et al. [10] was analyzed using the Seurat package [27]. The subset of CD4 T cells were selected based on the “free_annotation” property included in the dataset. Finally, the packages tidyverse [28], ggplot2, and qs were used for both flow cytometry and transcriptomic data.

### Statistics

All statistical analyses were performed using GraphPad Prism and were two-sided. Data was tested for normality with Shapiro-Wilk test and QQ-plot analysis. All results were obtained from at least two independent experiments and are shown as mean + SD. Statistical significance is defined as p<0.05 and shown as *, p<0.01 as **, p<0.001 as ***, and p<0.0001 as ****. p-values between 0.10 and 0.05 are indicated by the exact value.

## Acknowledgements

We would like to thank the Joller and Oxenius group members for helpful discussions. This work was supported by the Swiss National Science Foundation (310030_197590 to N.J.).

## Author contributions

Conceptualization: N.Y. and N.J. Experimentation: N.Y., E.G and C.P. Human Sample Collection: M.R. Analysis: N.Y. Formal Analysis: N.Y. Writing – Original Draft: N.Y. and N.J. Review and Editing: all authors. Funding Acquisition: N.J. Supervision: N.J.

## Supplementary information

There is no supplementary information for this paper.

